# Constraint-based metabolic control analysis for rational strain engineering

**DOI:** 10.1101/2020.11.26.399576

**Authors:** Sophia Tsouka, Meric Ataman, Tuure Hameri, Ljubisa Miskovic, Vassily Hatzimanikatis

## Abstract

The advancements in genome editing techniques over the past years have rekindled interest in rational metabolic engineering strategies. While Metabolic Control Analysis (MCA) is a well-established method for quantifying the effects of metabolic engineering interventions on flows in metabolic networks and metabolic concentrations, it fails to account for the physiological limitations of the cellular environment and metabolic engineering design constraints. We report here a constraint-based framework based on MCA, Network Response Analysis (NRA), for the rational genetic strain design that incorporates biologically relevant constraints, as well as genome editing restrictions. The NRA core constraints being similar to the ones of Flux Balance Analysis, allow it to be used for a wide range of optimization criteria and with various physiological constraints. We show how the parametrization and introduction of biological constraints enhance the NRA formulation compared to the classical MCA approach, and we demonstrate its features and its ability to generate multiple alternative optimal strategies given several user-defined boundaries and objectives. In summary, NRA is a sophisticated alternative to classical MCA for rational metabolic engineering that accommodates the incorporation of physiological data at metabolic flux, metabolite concentration, and enzyme expression levels.

## Introduction

Recent improvements in genome editing techniques have paved the way for more sophisticated and performant metabolic engineering designs for achieving desired physiological states of host organisms. Two approaches for reaching the targeted states exist: (i) integrating heterologous pathways to disruptively overcome native control patterns, and (ii) modifying the endogenous regulatory architecture by removal of the existing control loops (Bailey, 1991). The former method can be rather arduous because it requires testing if the integration of DNA fragments into the original genome sequence perturbs cellular regulation in the desired fashion. The latter technique demands knowledge about cellular control so that the DNA sequence can be modified effectively and without unwanted side effects.

Mathematical models are nowadays becoming an indispensable part of strain design. Available gene-protein-reaction associations of various organisms provide invaluable information about cellular metabolism and enable the elaboration of these models. The models can be studied computationally to interrogate and analyze cellular behavior and derive metabolic engineering strategies for improved cellular performance (Gombert and Nielsen, 2000). Strain design requires the identification and engineering of pathways toward the production of desired compounds (Hadadi and Hatzimanikatis, 2015), and mathematical models can provide an invaluable insight in the process of selection of deletions, insertions, and up- and down-regulation of genes encoding for metabolic enzymes. Reviews of the most prominent computational tools and workflows for the strain design are provided elsewhere (Costa et al., 2016; Long et al., 2015; Wang et al., 2017).

Metabolic control analysis (MCA) is a mathematical formalism that uses models to quantify the distribution of control over metabolic states in a network such as fluxes and concentrations (Kacser et al., 1995). In MCA, Control Coefficients (CCs) quantify how a given metabolic flux or metabolite concentration would respond to perturbations of the system parameters. This information is used in traditional rational metabolic design to identify the rate-limiting steps of the network and select potential targets for engineering. Strain engineering typically requires a holistic approach where one simultaneously analyzes the effects of genetic manipulations on specific productivity of desired molecules, maximum achievable yield, energetic and redox requirements, etc. Simultaneous analysis of these effects is a cumbersome task using classical MCA tools, especially if the design requires multiple genetic alterations. Moreover, MCA does not allow including explicitly any form of physiological or design constraints, which can lead to unrealistic predictions.

We present here Network Response Analysis (NRA), a constraint-based workflow that aims to tackle these obstacles. NRA utilizes populations of CCs to consistently derive metabolic engineering strategies and trace the effects of multiple parameter perturbations. The advantage of this method is that physiologically relevant bounds and constraints can be imposed to the system, as opposed to the classical MCA. NRA is inspired by the work by Hatzimanikatis et al. (1996a); (1996b) who proposed a Mixed Integer Linear Programming (MILP) formulation for querying cellular responses upon enzymatic perturbations that uses MCA-based flux and concentration CCs. Therein, the authors applied their formulation on simple linear and branched pathways to propose metabolic engineering strategies. Here, we extend this formulation to allow for studying larger scale metabolic systems with guarantied thermodynamic feasibility.

To illustrate how NRA can be used to efficiently analyze, enumerate, and propose alternative metabolic engineering strategies, we used a large-scale thermodynamically-curated, metabolic model of *E. coli* (Hameri et al., 2019c), which describes the central carbon pathways in aerobic growth conditions. Using the stoichiometric model as a scaffold, we employed the ORACLE framework (Andreozzi et al., 2016a; Chakrabarti et al., 2013; Hameri et al., 2019b; Miskovic et al., 2017; Miskovic and Hatzimanikatis, 2010; Soh et al., 2012; Tokic et al., 2020) to generate populations flux and concentration CCs consistent with the experimental observations. We then used the generated CCs to formulate with NRA the design strategies in two case studies (i) improvement of glucose uptake rate, and (ii) maximization of specific productivity rate of pyruvate while preserving a pre-specified yield of pyruvate from glucose. These studies clearly show the potential, flexibility, and ease of use of NRA when realistic, multi-objective requirements for the strain design should be met.

## Results and Discussion

### NRA method

The first step of the NRA method is the selection and curation of a metabolic network that captures the physiology of a studied organism (Fig. 1). Then, we calculate the relevant flux and concentration CCs (FCCs and CCCs) that describe the network’s responses to parameter perturbations such as modifications of enzymatic activities with the ORACLE framework, which makes use of Monte Carlo sampling (Miskovic and Hatzimanikatis, 2011; Wang et al., 2004). Finally, we use the computed sets of CCs along with the user-defined requirements and additional physiological constraints to construct a constraint-based MILP optimization problem (Fig. 1). The user-defined inputs depend on the studied problem and design limitations, and these typically include the number of desired gene manipulations, minimal allowable specific productivity, minimum allowable yield, etc. From experimental measurements or assumptions on physiology, we can infer physiological constraints such as allowable (or desired) bounds on fluxes and concentrations in the metabolic network.

**Figure 1.**
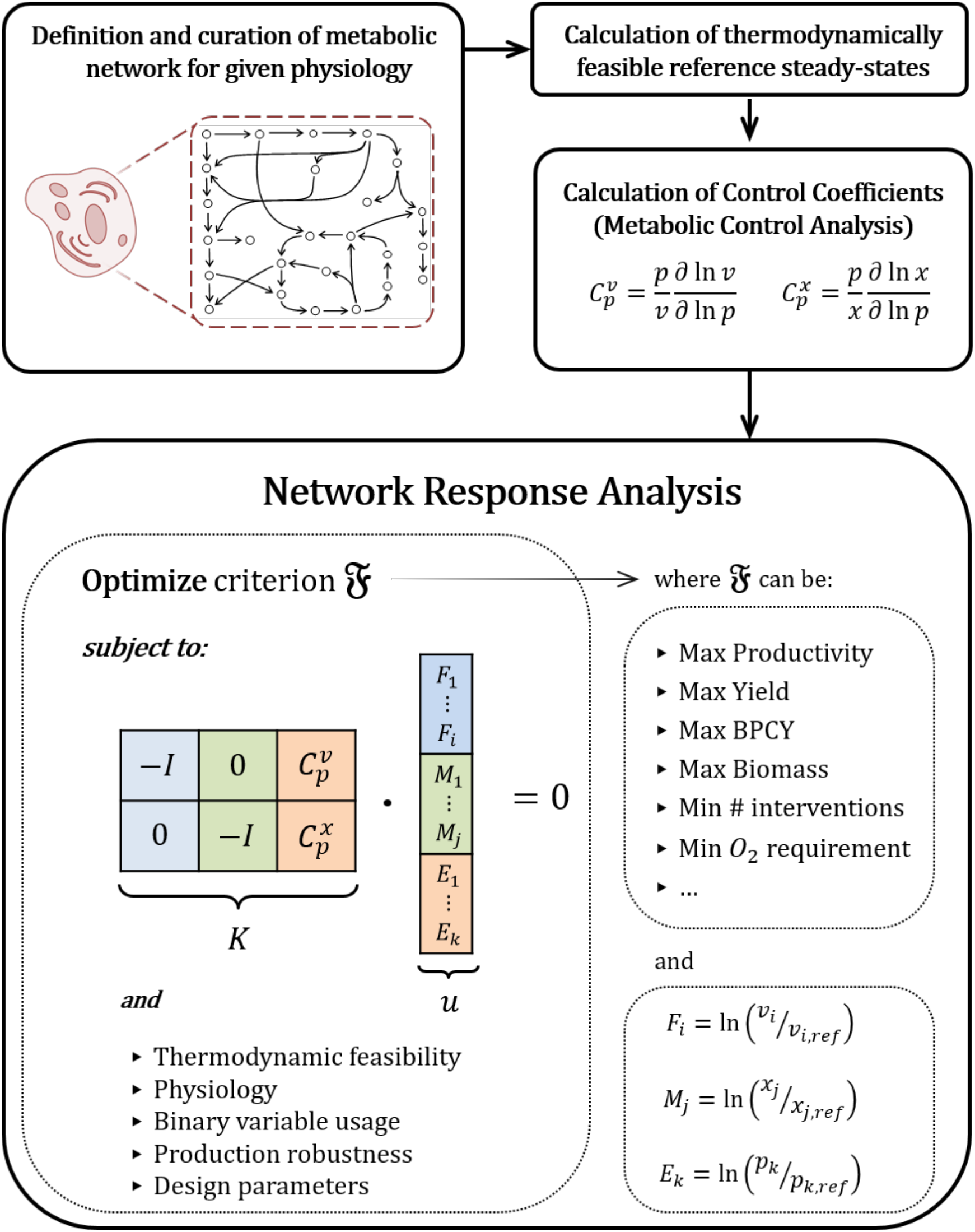
The NRA workflow is organized in four main steps. In the first three steps, we formulate the stoichiometry, integrate available experimental data and compute the steady-state thermodynamically feasible fluxes and concentrations, and compute the flux and concentration control coefficients for the studied physiological condition. In the fourth step, metabolic engineering strategies are devised by solving a MILP. Criterion 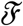 and additional constraints can be chosen from a set of metabolic engineering criteria such as the ones provided in Table 1. Variables F_i_, M_j_ and E_k_ are thelogarithmic deviations in flux, metabolite concentration and parameter with respect to their respective reference steady states (Eq. 23), and their bounds define the solution space of the optimization problem (Eqs. 8-10). The definition of the other optimization variables and parameters is given in Table 2.

The outcome of the NRA optimization are sets of alternative combinations of genes that should be engineered to improve the cellular performance given the imposed user-defined inputs and physiological constraints. A principal advantage of the MILP formulation is that it allows the user to introduce constraints on metabolic states and additional relevant design constraints to the system, thus simultaneously offering flexibility and tight control over the rational strain design.

### NRA form ulation

The NRA core equations can be expressed in a matrix-vector form (Table 1, Eq. 7) similar to the ones of Flux Balance Analysis (FBA) (Orth et al., 2010) and Thermodynamics-based Flux Analysis (TFA) (Henry et al., 2007; Salvy et al., 2019). NRA accommodates a wide gamut of design objectives, such as the maximization of productivity or product yield (Eqs. 1-2), biomass-product coupled yield (BPCY) (Eq. 3), the maximization of biomass formation (Eq. 4), the minimization of required genome-editing interventions (Eq. 5), and the minimization of oxygen requirements (Eq. 6) (Klamt et al., 2018; Patil et al., 2005; Schneider and Klamt, 2019; Varma et al., 1993). Since we have defined the NRA variables in logarithmic form, we can express the otherwise nonlinear objectives like yield or BPCY in a linear form, rendering the solution of the mathematical problem easier to attain than with formulations such as FBA.

**Table 1.**
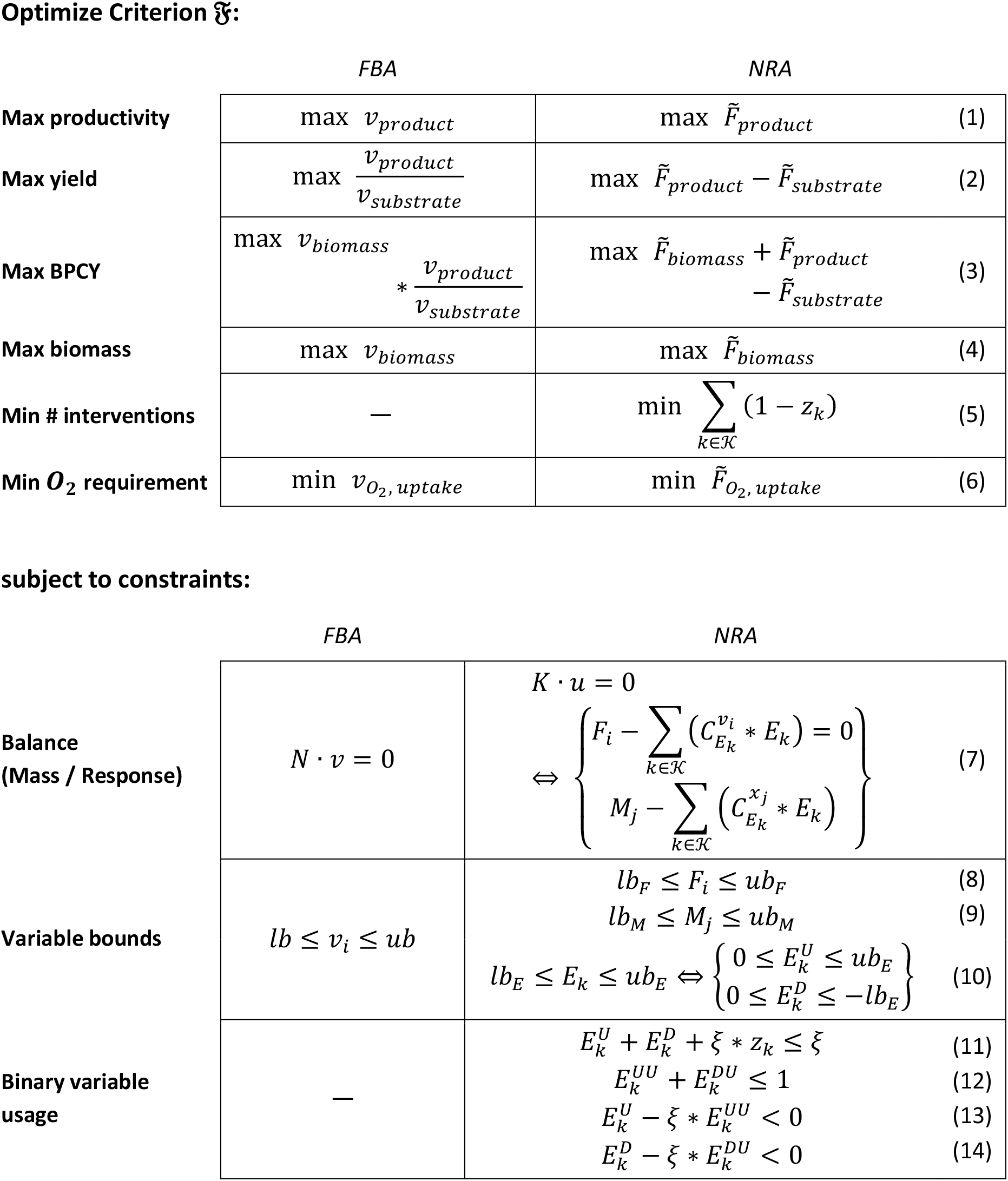
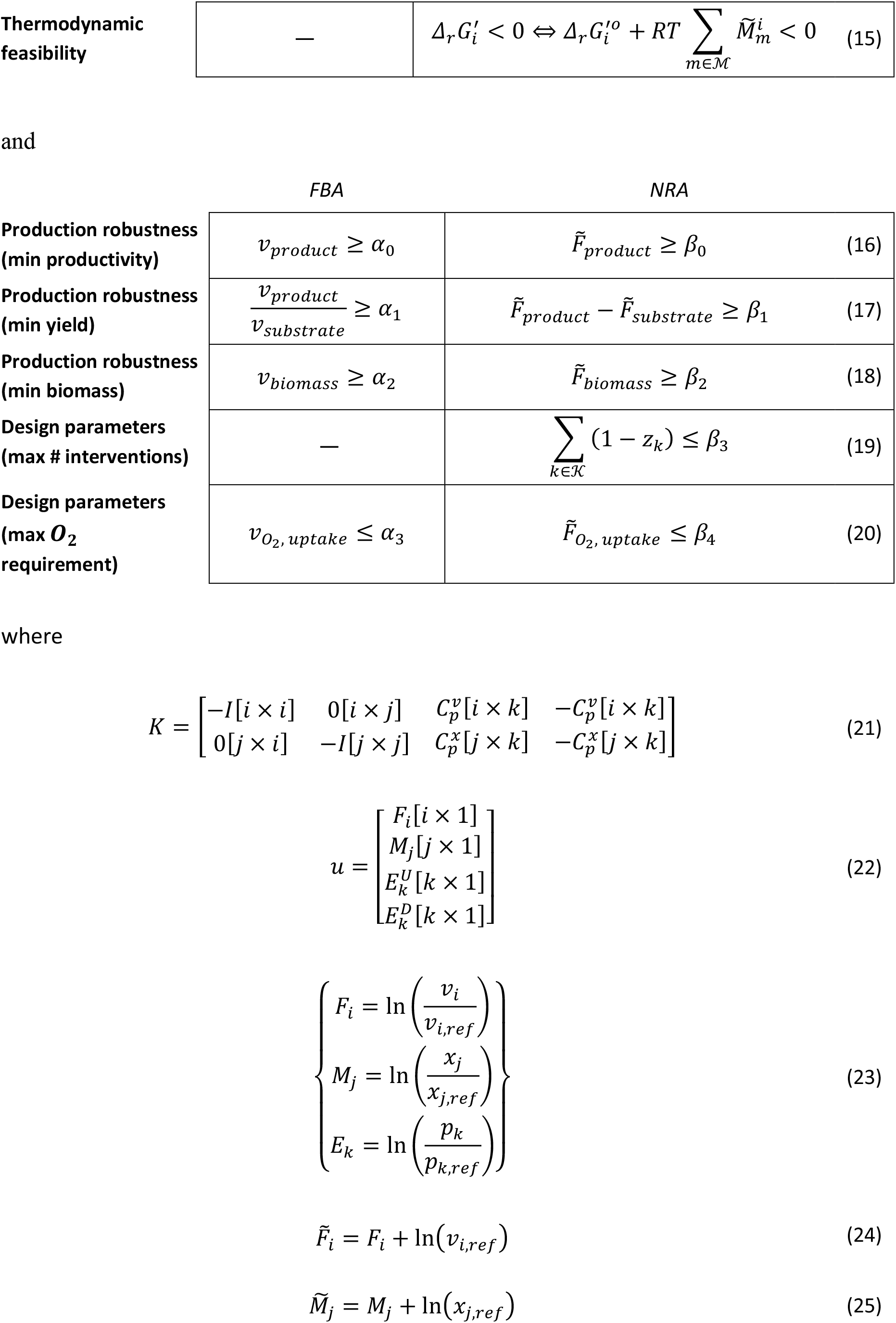
The NRA mathematical formulation together with a non-exhaustive selection of optimization objectives. The definition of indices, parameters, and variables is provided in Table 2.

Importantly, the NRA formulation allows us to prevent thermodynamically infeasible designs because it naturally includes thermodynamic constraints regarding the Gibbs free energy change 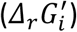 of each reaction (eq. 15). Furthermore, the proposed formulation allows imposing additional design criteria such as production robustness and operational parameters (Eqs. 16-19). The NRA optimization problems can be solved with the TFA toolbox (Salvy et al., 2019). We provide more details about the NRA formulation in Methods.

### Strain design with physiological and design constraints for improved glucose uptake

Metabolic engineering interventions on pathways inevitably result in altered reaction rates as well as metabolite concentration levels. NRA, being a constraint-based method, allows for setting appropriate constraints on these quantities. Both fluxes and concentrations need to be constrained within realistic physiological bounds, conditional to each case study. For instance, severe changes in metabolic concentrations upon metabolic engineering interventions could significantly influence the organism’s growth or even lead to an excess of toxic byproducts. The strain design should likewise consider that enzyme expression levels cannot increase beyond the currently reported experimentally achievable levels, and it cannot allow an infinite increase of reaction fluxes in the network. In contrast, the design should also be able to model gene knockouts by allowing both enzyme activities and reaction fluxes to decrease close to zero.

Here, we examined the effects of the imposed physiological and design constraints on the strain design for improved glucose uptake. To this end, we analyzed the achievable glucose uptake rates with 2-fold, 5-fold, and 10-fold maximum allowable deviation of enzyme activities from the reference level for a set of designs ranging from 1 to 25 gene manipulations (Figure 2a). The metabolite concentrations were subject to the thermodynamic feasibility constraints (Methods), and within the predefined physiological ranges (10nM – 0.1M) for each cellular compartment. We allowed the fluxes to increase up to 10-fold of their reference level, and both fluxes and enzyme activities could reduce to zero. The latter means that solutions can include potential gene knockouts. As a mean to investigate the sensitivity of obtained solutions, we repeated the study for one reference and 18 extreme sets of control coefficients (Methods).

**Figure 2.**
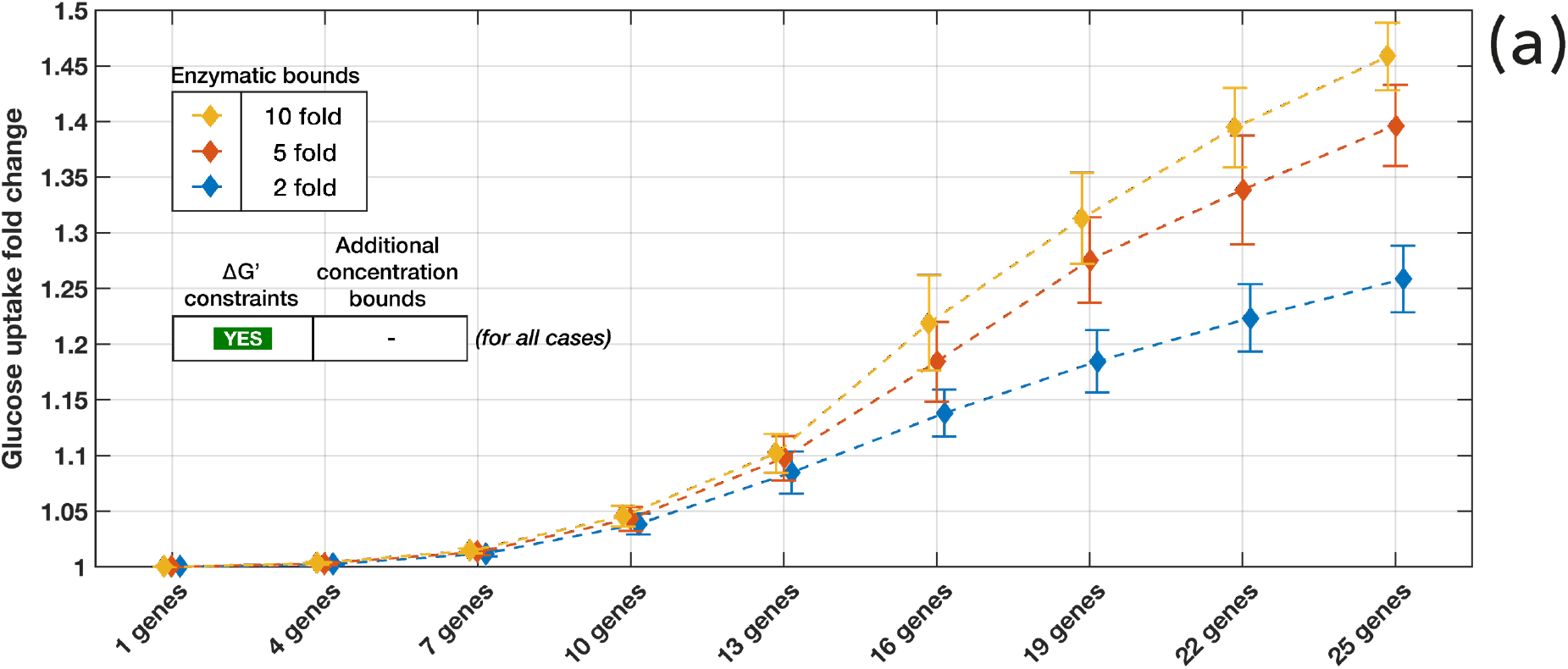
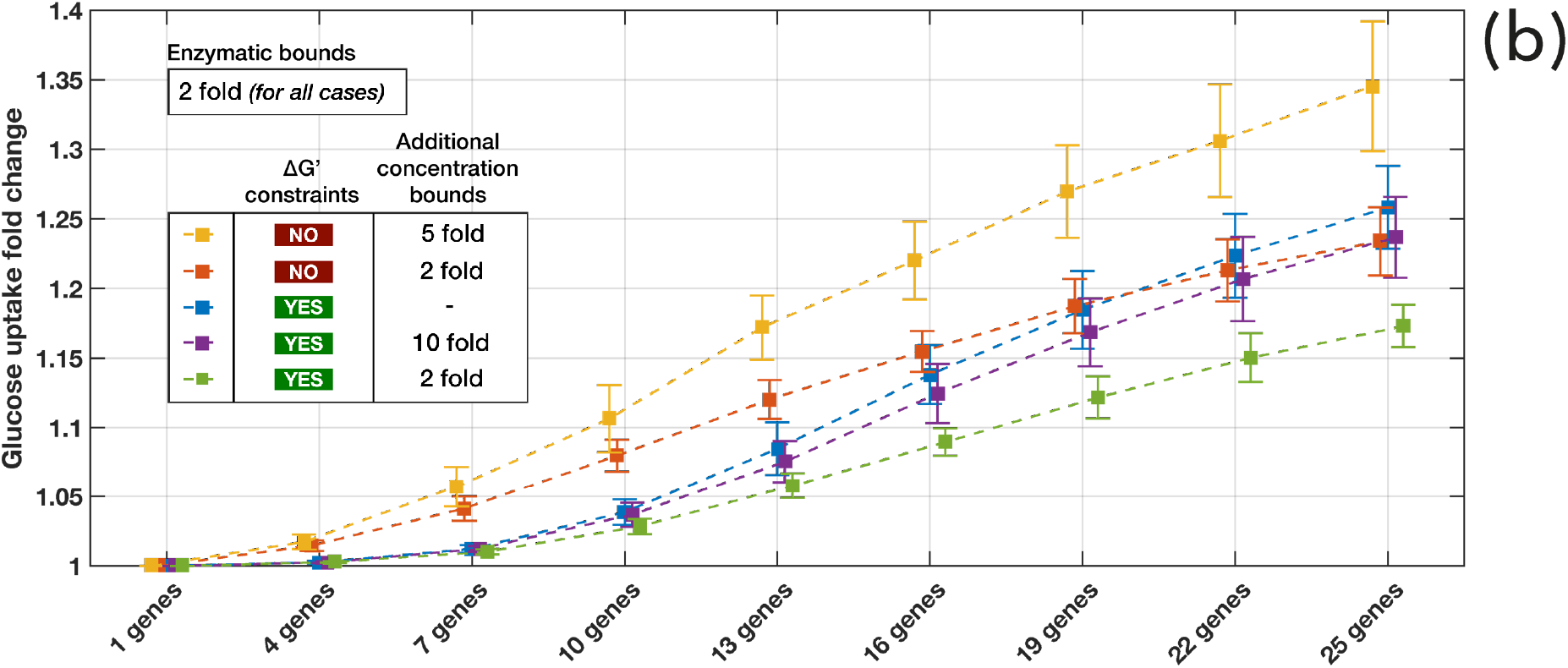
*Effects of the physiological and design constraints on glucose uptake rate for a set of designs with different number of gene manipulations. Effects of: (a) allowed 2-fold (blue), 5-fold (orange), and 10-fold (yellow) changes in enzyme perturbation magnitude, and (b) different imposed metabolite concentration bounds. The study was performed for one reference and 18 extreme sets of CCs selected using PCA (Methods). In all cases, the fluxes were allowed to increase 10-fold and decrease to zero.* The whiskers and the respective symbols indicate the interquartile ranges and the means of the considered CC-sets, respectively, as adjusted by the Bonferroni *correction (Methods). Blue lines correspond in both graphs.*

As the allowable enzyme activity change (Eqs. 10 and 23) increased from 2-to 10-fold, the predicted attainable glucose uptake rate was about the same for up to 10 gene manipulations, indicating that for a small number of gene manipulations the upper limits on enzyme activity were not a limiting factor (Figure 2a). However, starting from 13 gene manipulations, the difference between the predictions increased considerably. As expected, the higher limits on enzyme activity, the larger predicted improvement of glucose uptake was observed. For example, NRA predicted for 25 gene manipulations that glucose uptake rate would increase by 26%, 39%, and 46% for 2-, 5-, and 10-fold change in enzyme activity, respectively. Interestingly, the predicted fold change of the glucose uptake across the nineteen studied reference and extreme CC-sets varied similarly for the designs with 13 or more gene manipulations (Figure 2a whiskers). This rather constant variability as we go toward a higher number of gene manipulations suggests that variability among 19 sets is primarily determined by the activity of a relatively small number of enzymes, which predominantly have control over the glucose uptake rate. This finding is in line with previous studies of metabolic systems demonstrating that just a few enzymes in the network (or corresponding parameters) determine the key metabolic properties such as system stability (Andreozzi et al., 2016b) or control over production fluxes (Miskovic et al., 2019a). A similar observation was reported in a more general context of biological systems (Daniels et al., 2008; Gutenkunst et al., 2007).

Next, we investigated how constraints on concentration deviations (Eqs. 9 and 23) affect the attainable glucose uptake. This is a salient aspect of strain design because metabolic engineers have to ensure that metabolite concentrations remain within physiological bounds. For instance, it is vital not to exceed toxicity levels for some compounds. The studies on the effects of metabolite concentration constraints have also to consider thermodynamics because it is well known that the standard free Gibbs energy change of reactions couples the reaction directionalities and the metabolite concentrations (Ataman and Hatzimanikatis, 2015). For this analysis, we have performed several studies by imposing different concentration bounds together with and without thermodynamic constraints (Figure 2b). In general, our results suggest that NRA without thermodynamic constraints tends to overpredict the increase in glucose uptake (Figure 2b), meaning that thermodynamic constraints are limiting factors of strain design. The notable exception was that, starting from 19 gene manipulations, the 2-fold constraints on concentrations are more limiting than the thermodynamic ones (Figure 2b blue & orange lines). As expected, our results also show that the tighter the concentration deviation bounds we impose, the less important improvements of glucose uptake could be attained (Figure 2b). For example, the attainable increase of glucose uptake rate with the thermodynamic and additional 2-fold and 10-fold constraints for 25 gene manipulations were 17% and 24%, respectively (Figure 2b, green and violet). We also observed that the variance of glucose uptake increase was smaller as the concentration bounds became more constrained. Similarly, we observed a trend that the variance in the studies with the thermodynamic constraints was smaller than in the ones without thermodynamic constraints.

### Metabolite concentrations limiting the glucose uptake

Having demonstrated that limits on metabolite concentrations, either thermodynamic constraints or physiological limitations, significantly affect the attainable glucose uptake, we investigated how many and which metabolite concentrations should violate the thermodynamic constraints to achieve a higher glucose uptake. For simplicity and clarity of exposition, we allowed designs with one, two, four, and seven gene manipulations (Figure 3). In the cases of one and two gene manipulations, the flux through glucose uptake could not be modified with the thermodynamically feasible concentrations (zero violations). For a larger number of gene manipulations, a small increase in glucose uptake could be achieved even without violating the thermodynamics. For example, the manipulation of seven genes would yield ~2% of glucose uptake increase for zero violations. However, when we allowed some concentration deviations to exceed their bounds, the potential violations pushed the attainable glucose uptake to higher values (Figure 3). For instance, the seven gene manipulations design with ten concentration violations would result in 5.5% increase in glucose uptake.

**Figure 3.**
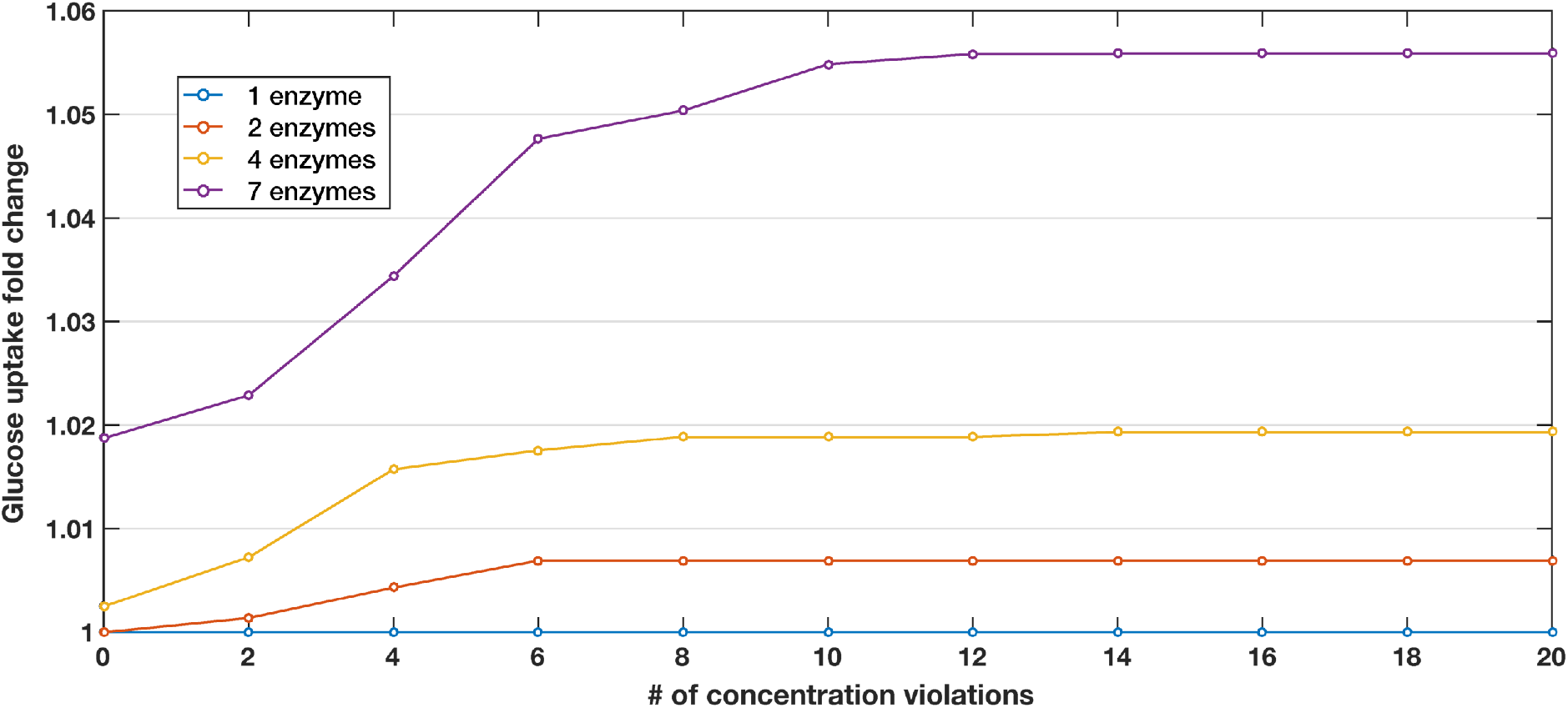
Maximal attainable increase in glucose uptake as a function of a different number of metabolite concentration violations for one, two, four, and seven gene modifications. The fluxes were allowed to increase 10-fold and decrease to zero, the non-violated concentration bounds were subject to the thermodynamic constraints, and the enzymatic bounds were set to 2-fold. The reference model was used for all cases.

Next, we focused on finding which were the metabolites whose concentration constraints should be violated to improve glucose uptake. To this end, we studied the case of four violations and two, four, and seven gene manipulations. For each gene manipulation study, we obtained the unique sets of four metabolite concentrations violating constraints (Table 3a). The three gene manipulation studies involved, in total, seven species with concentrations violating the thermodynamic constraints. Among the seven species, peroxisomal protons appeared in all three studies. Moreover, irrespectively of the study, to achieve a higher glucose uptake, the concentrations of protons (both cytosolic and peroxisomal), AMP, and phenylalanine needed to be increased, while the ones for CTP, dCTP and glutamine needed to be decreased. The violations ranged from 2% for the case of cytosolic hydrogen to 57% for the case of CTP (Supplementary Table S1).

**Table 2.**
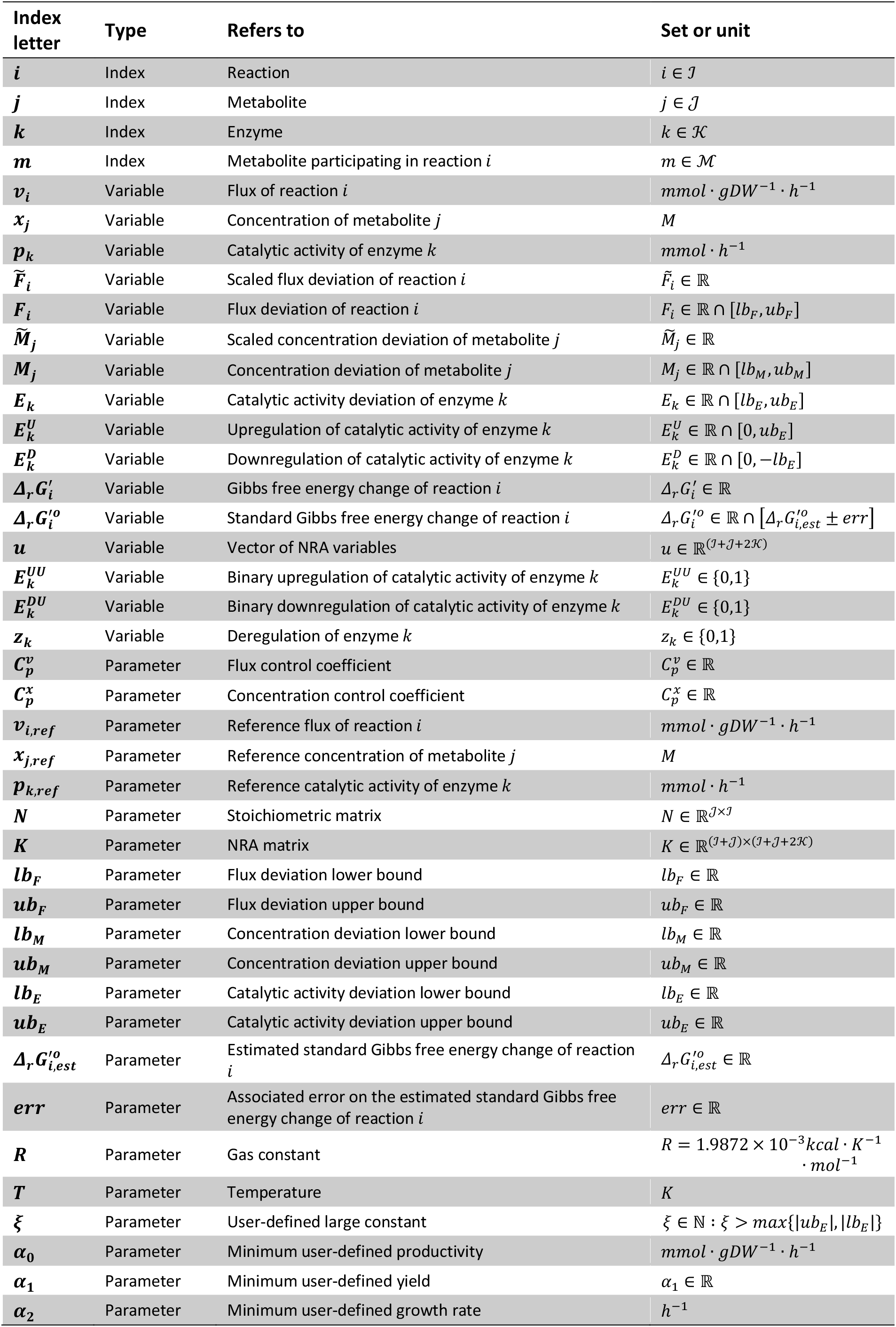
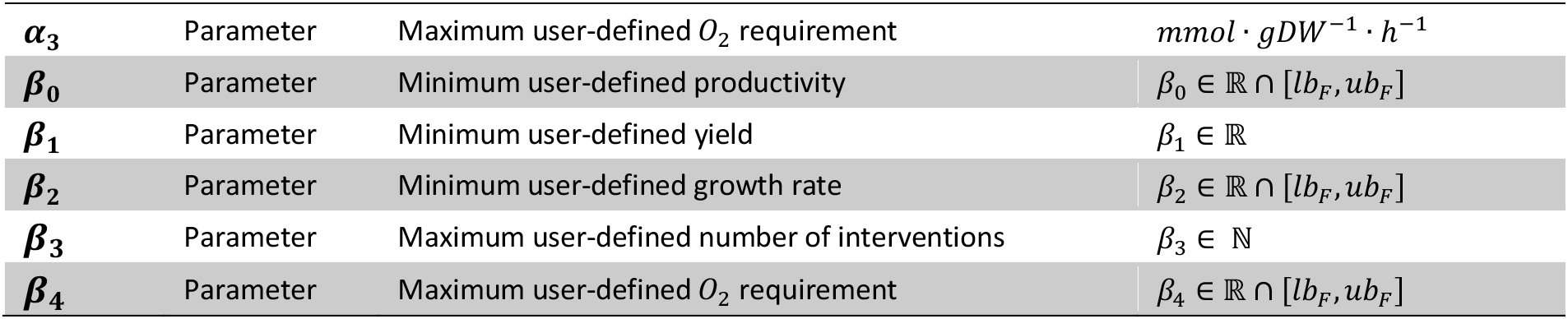
Indices, variables, and parameters used in the NRA formulation.

**Table 3.** Sets of four metabolite species with concentrations violating thermodynamic constraints for designs with two, four, and seven gene manipulations. The arrows indicate should a metabolite concentration be increased or decreased to improve glucose uptake. c: cytosol, p: periplasm.

|  |  | # of parameter manipulations |  |  |
| --- | --- | --- | --- | --- |
|  |  | 2 | 4 | 7 |
| Metabolite concentrations | AMP (c) ↑ | X | - | X |
|  | CTP (c) ↓ | - | X | X |
|  | dCTP (c) ↓ | - | X | - |
|  | H<sup>+</sup> (c) ↑ | X | X | - |
|  | H<sup>+</sup> (p) ↑ | X | X | X |
|  | L-Glutamine (c) ↓ | - | - | X |
|  | L-Phenylalanine (c) ↑ | X | - | - |

This analysis provides an opportunity to focus on each of the identified molecules, draw hypothesis about their role in the system limitations, and investigate these interplays and ways to overcome them *in vitro.* For example, it suggested that the pH value in compartments can be a limiting factor for metabolic design.

### NRA design for Pyruvate production considers together specific production rate and yield

Pyruvate (pyruvic acid) is widely used in the food, chemical, and pharmaceutical industries. It is a precursor for the synthesis of various amino acids, and has been used for the production of antioxidants, food additives and supplements, pharmaceutical precursors, and biofuels (Atsumi et al., 2008; Kalman et al., 1999; Li et al., 2001; Zhang et al., 2010). The microbial production of pyruvate has been largely explored, and has involved both strain and process engineering and development (Maleki and Eiteman, 2017). In *E. coli,* pyruvate has been identified as one of the main hubs for the production of non-native commercial products (Zhang et al., 2016). The most common approach in microbial engineering for the overproduction of pyruvate is through deletions of the downstream utilization of pyruvate towards byproducts such as acetate, acetyl-CoA, and ethanol among others (Akita et al., 2016; Causey et al., 2004; Zhu et al., 2008).

To illustrate the features and flexibility of the NRA method, we showcase design for the improved specific productivity rate of pyruvate while taking into account the yield of pyruvate from glucose, design constraints, and thermodynamic feasibility. We imposed the following design and physiology constraints: (i) up to five gene/enzyme activity manipulations, (ii) the genes encoding for metabolic enzymes could either be upregulated up to 50-fold or downregulated down to a knockout, (iii) the fluxes could increase up to 100-fold for upregulation and decrease down to zero for knockouts, and (iv) the concentration values were subject to the thermodynamic feasibility constraints and the physiological ranges (10nM – 0.1M). Given these constraints, we first performed an optimization to determine the maximum yield of pyruvate from glucose. Then, we added the pyruvate yield to be at least 90% of this value to the set of constraints and maximized the specific pyruvate productivity rate. In this manner, we were able to implicitly account for the potential tradeoffs of yield and productivity that can occur in such designs.

We generated 51 alternative designs with five gene manipulations providing at least 99% of the maximum specific productivity rate of pyruvate and fulfilling the imposed constraints. The alternative designs involved the manipulation of genes corresponding to 48 distinct enzymes (Supplementary Table S2). All cases provided over a 22-fold increase in both the pyruvate yield and specific productivity rate compared to the reference state. To understand better the mechanisms and identify metabolic patterns behind improved pyruvate production and yield, we performed clustering analysis over 51 designs with respect to (i) the 48 enzyme activity manipulations (Figure 4), and (ii) predicted change in metabolic fluxes upon changes in enzyme activities (Figure 5). For the clustering based on the absolute change in fluxes, we used the set of 67 reactions that had an absolute flux change of more than 0.01 mmol/gDW/h.

**Figure 4.**
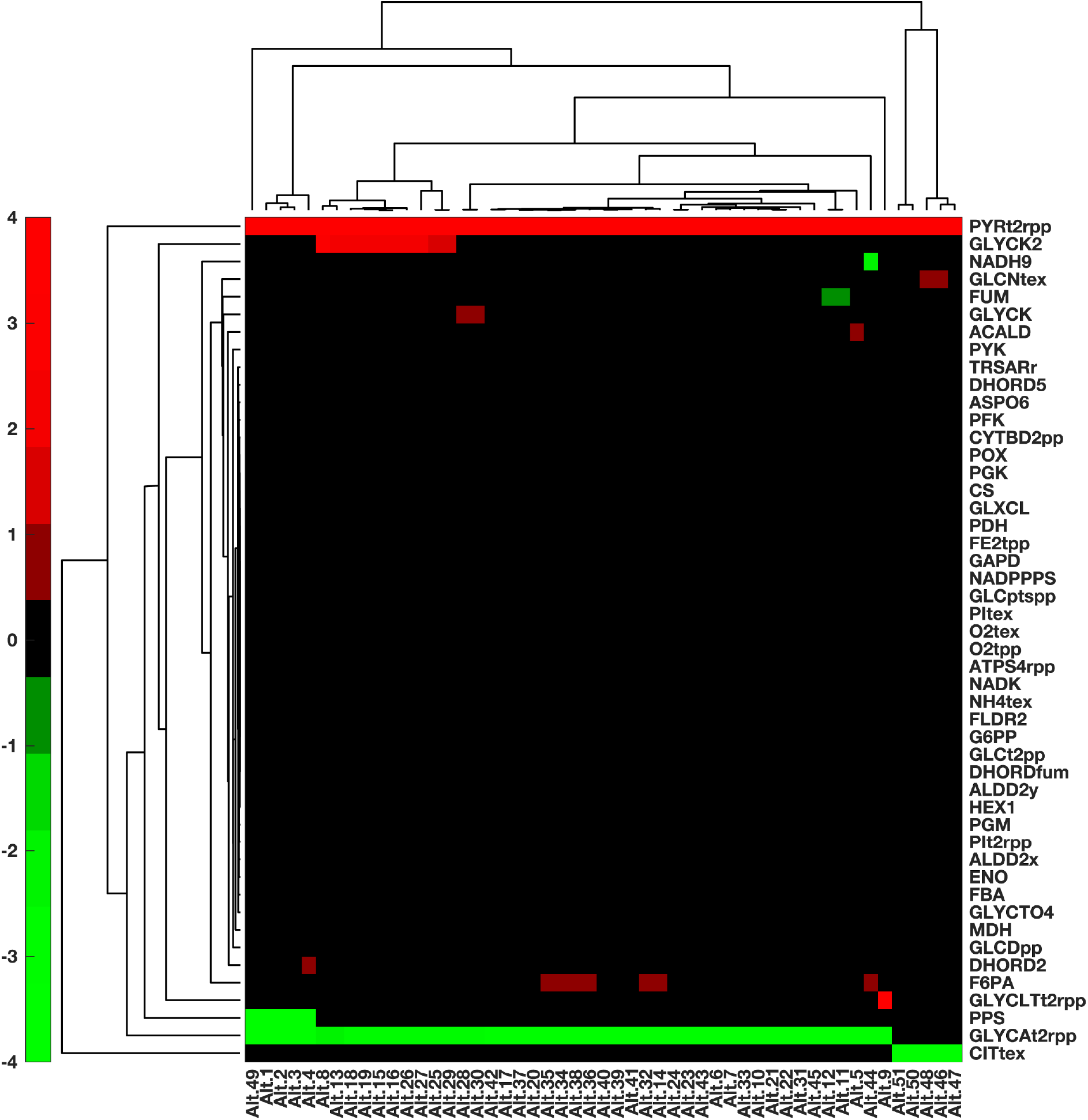
Hierarchical clustering of the 51 alternative designs for the increase of pyruvate productivity, based on the suggested deregulation of individual enzymes.

The transport of pyruvate from the cytosol to the periplasm (PYRt2rpp) appeared as a target in all designs with 50-fold upregulation of the PYRt2rpp encoding gene (Figure 4). The upregulation of glycolytic enzymes and enzymes leading to pyruvate synthesis would also improve pyruvate production, with the most prominent target being glycerate kinase (GLYCK2). We also observed knockouts (or significant downregulations) with the majority of downregulated genes involving the consumption of pyruvate towards the formation of byproducts. Among these, the periplasmic transport of glycerate (GLYCAt2rpp) was present in most generated sets, being replaced by the extracellular transport of citrate (CITtex) in a few cases (Figure 4). We also observed the knockout of PPS (Phosphoenolpyruvate synthase), which is associated with the conversion of pyruvate to phosphoenolpyruvate.

A closer cross-inspection of the two figures together with the Supplementary Table S2 reveals that there are five groups of alternative ways to satisfy design specifications. Alternatives 1-4, 20, 32, 34, 35, 37, 38, 44, and 49 (Figures 4 and 5, Supplementary Table S2) constituted the first group that improves pyruvate production while maintaining at least 90% of the yield by: (i) a strong upregulation of pyruvate transport PYRt2rpp; (ii) a strong downregulation of GLYCAt2rpp; and (iii) a slight upregulation of glycolysis either via enolase (ENO) for Alternative 49 or via fructose 6-phosphate aldolase (F6PA) for other alternatives in this group (Supplementary Table S2); (iv) a knockout of PPS for alternatives 1-4, 49 or a slight downregulation of fructose-bisphosphate aldolase (FBA) for alternatives 20, 32, 34, 35, 37, 38, and 44.

**Figure 5.**
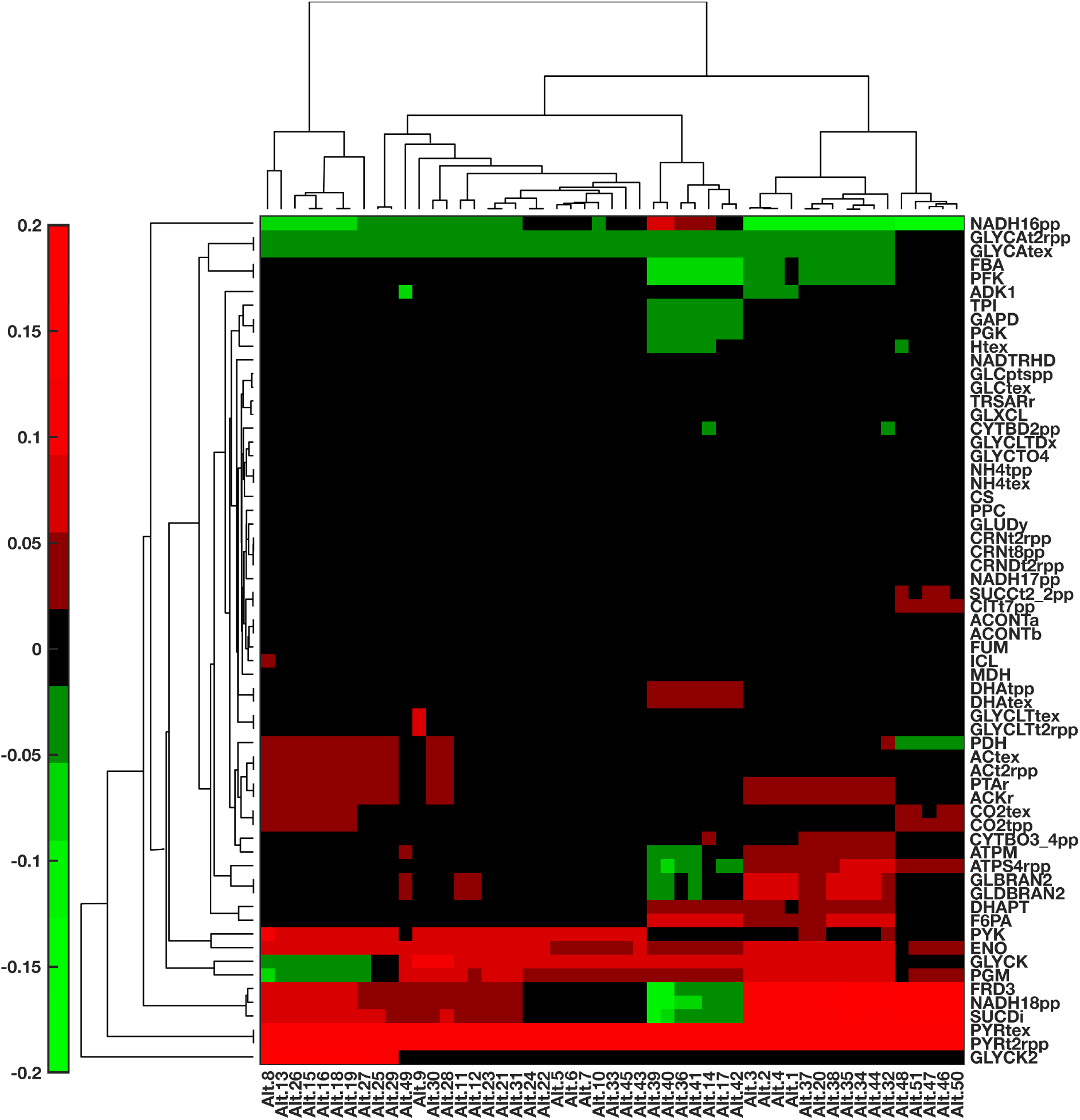
Hierarchical clustering of the 51 alternative designs for the increase of pyruvate productivity, based on the absolute change in flux value of the 67 most affected reactions in the network.

As a result of these manipulations, the carbon flow was re-directed from the secretion of (R)- glycerate toward the production of phosphoenolpyruvate through glycerate kinase (GLYCK), phosphoglycerate mutase (PGM), and ENO (Figure 6 and Supplementary Figure S1). Downstream, phosphoenolpyruvate is converted to pyruvate through dihydroxyacetone phosphotransferase (DHAPT), whose activity was also increased. This group is further characterized by a slight increase in acetate production and CO_2_ secretion, and a deregulation of the ATP metabolism such as an increase of the ATP non-growth associated maintenance (ATPM) or a decrease in activity of adenylate kinase (ADK1) for alternatives 1-4, 49. Moreover, the conversion of fructose-6-phosphate to glyceraldehyde-3-phosphate instead through FBA and phosphofructokinase (PFK) was diverted through F6PA.

**Figure 6.**
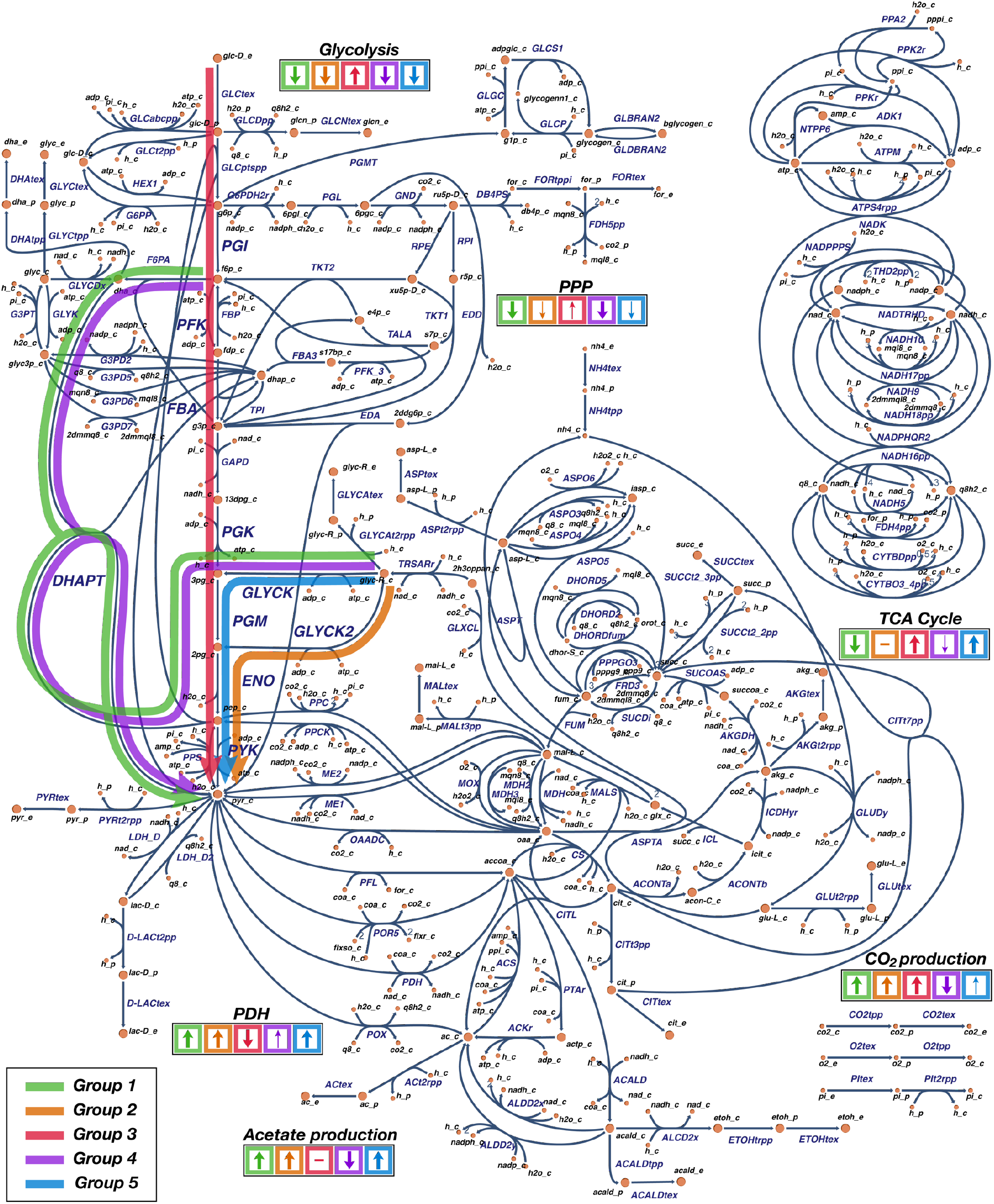
Overview of metabolic engineering strategies devised using NRA for the improved specific production rate of pyruvate while preserving the prespecified yield of pyruvate from glucose. 51 strategies devised with NRA were categorized in 5 distinct groups. The thick arrows on the graph denote the principal ways of carbon re-direction from the wild-type strain steady-state fluxes. The arrows in the colored boxes denote if the activity of the corresponding metabolic subsystem (glycolysis, pentose phosphate pathway (PPP), TCA cycle, acetate production, and CO_2_ production) or reaction (PDH) was increased (arrow up), decreased (arrow down), or remained unchanged (dash). The thicker arrow in the colored boxes, the higher change in the activity occurred.

The second group consisting of alternatives 8, 13, 15, 16, 18, 19, 25-27, and 29 shared the manipulations (i) and (ii) with the first group. In addition, this group involved: (iii) an upregulation of glycerate kinase GLYCK2; and (iv) a slight upregulation of pyruvate kinase (PYK). The observed effects of these manipulations were similar to the ones of the first group with the increased activity of lower glycolysis and acetate secretion pathway (Figures 5, 6 and Supplementary figure S2). The notable difference was that the carbon diverted from glycerate secretion was channeled through GLYCK2, ENO, and PYK to pyruvate. Furthermore, we observed a slight increase in activity of the TCA cycle and pyruvate dehydrogenase (PDH), whereas the ATP metabolism remained mostly unchanged.

The third group formed by alternatives 46-48, 50, and 51 was distinct from the other groups because it involved strategy to knockout citrate transport CITtex instead of GLYCAt2rpp (Figure 4). Additional manipulations in these group were a slight downregulation of citrate synthase (CS) and a slight upregulation of glycolytic enzymes PGM (Alternatives 46, 47, 50, 51) or ENO (Alternative 48). Overall, these manipulations resulted in increased activity of the upper and lower glycolysis, pentose phosphate pathway, and the TCA cycle (Figure 6 and Supplementary Figure S3). This was the only group with increased activity of the upper glycolysis. We have also observed a decrease in activity of PDH (Figures 5 and Supplementary Figure S3).

The fourth group constituted by alternatives 14, 17, 36, 39-42 had a distinct pattern in the network flux distributions while sharing manipulations (i)-(iii) with the first group (Figure 5, 6 and Supplementary Figure S4). A slight downregulation of PFK together with manipulations (i)-(iii) had a considerable impact by reducing the activity of the reactions in the upper glycolysis (PFK, FBA, triose-phosphate isomerase (TPI), glyceraldehyde-3-phosphate dehydrogenase (GAPD), phosphoglycerate kinase(PGK)), the ETC chain (NADH dehydrogenase (NADH18pp), Cytochrome oxidase bo3 (CYTBO3_4pp) and the ATP metabolism (ATPM and ATP synthase (ATPS4rpp)). We have also observed, in contrast to other groups, a reduced activity in CO_2_ and acetate secretion pathways.

The fifth group, composed of alternatives 5-7, 9-12, 21-24, 28, 30, 31, 33, 43, 45, and the second group have in common manipulations (i), (ii), and (iv) (Figure 4 and Supplementary Table S2). Additionally, the fifth group involved either a very slight upregulation of glycolytic enzymes PGM, ENO, and PGK (alternatives 11, 12, 21-24, 31, 43, and 45) or a very slight downregulation of PDH (alternatives 5-7, 10, 28, and 30). As expected, the resulting flux distribution was similar to the one of the second group (Supplementary Figure S5). The difference was that in this group the carbon from (R)-glycerate was diverted to 2-phosphoglycolate through GLYCK and PGM instead through GLYCK2 as it was done in the second group. Overall, compared to other groups, the manipulations of this group have changed the least the network flux distribution (Figure 5).

Once the principal strategies are determined, the final decision is made by experts based on the comparative analysis of the proposed alternative groups and on considerations about the practical implementation of the designs.

### Comparison with targets determined by looking only at unconstrained specific productivity

We proceeded by examining how different are the targets obtained with the NRA design from the ones determined by looking only the specific productivity rate of pyruvate. This comparison will reveal how the physiology and design constraints affect our design decisions. To this end, we computed the mean values of the control coefficient of the specific productivity rate of pyruvate with respect to network enzyme activities, and then ranked them according to their absolute value. Most of the top 15 enzymes represent either extracellular transports such as oxygen uptake and ammonium secretion, as well as glycolysis reactions leading to the synthesis of pyruvate (Table 4). Interestingly, the majority of these enzymes do not appear as targets in any of the NRA alternatives (Table 4 and Supplementary Table S2). Some of these enzymes exhibit a large control over multiple fluxes and concentrations across the metabolic network. These are, therefore, severely constrained by the imposed specifications in the constrained NRA design. This suggests that the NRA formulation will favor parameters that have less control over the network, ensuring that cellular balance will not be excessively perturbed.

**Table 4.**
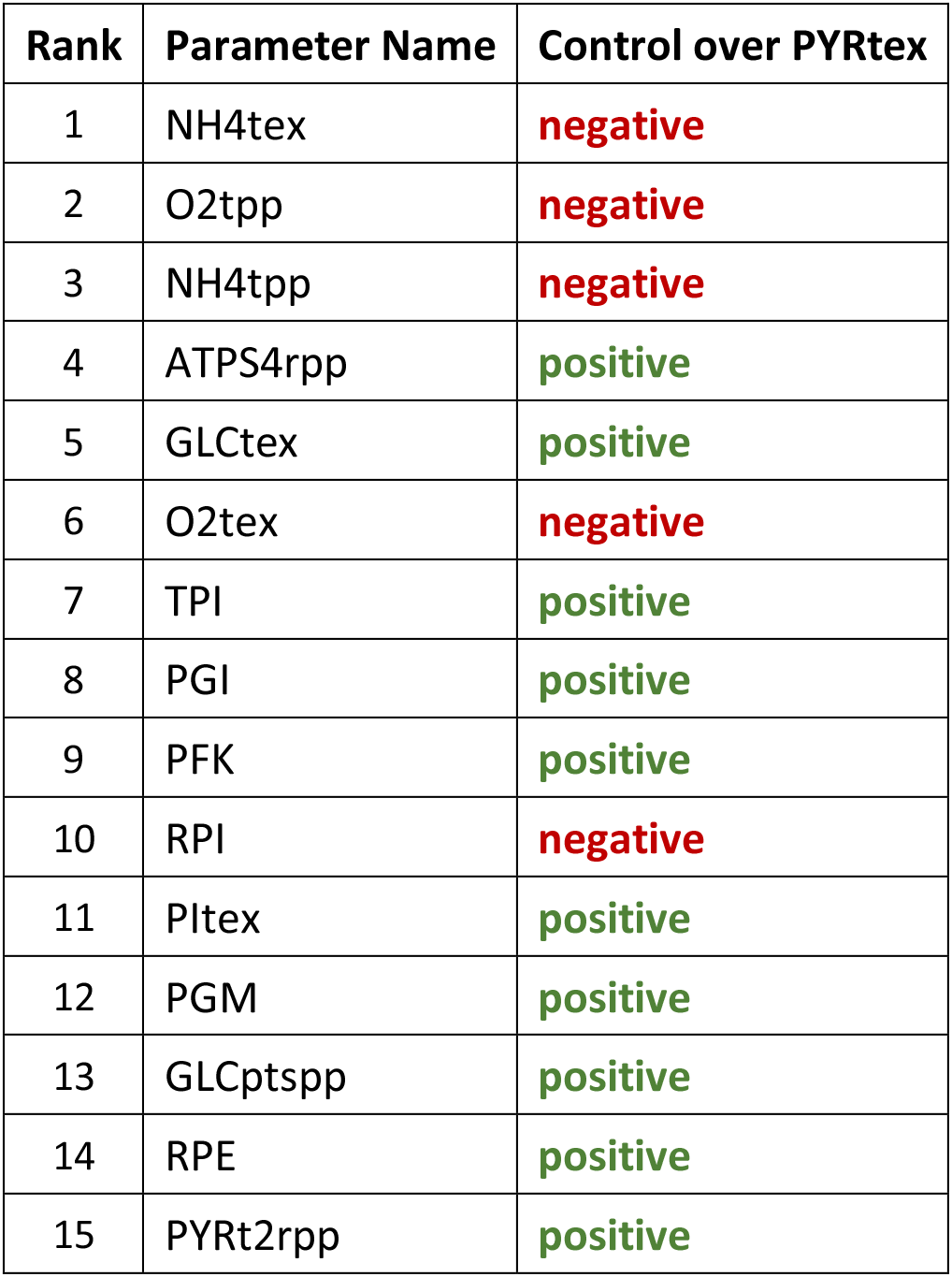
Top ranked parameters based on their control over pyruvate production flux PYRtex. Ranking was computed based on the mean values of 50’000 sets of Control Coefficients.

## Materials and Methods

### Metabolic Control Analysis notions

In MCA, the CCCs, 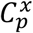, and the FCCs, 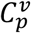, are defined as the fractional change of metabolite concentrations *x* and metabolic fluxes *v*, respectively, in response to a fractional change of system parameters *p* (Hatzimanikatis and Bailey, 1996; Kacser et al., 1995). These CCs serve as measurable outputs that provide information about the levels of control that system parameters have on the studied biological system and physiology. From the log(linear) formalism (Hatzimanikatis et al., 1996a; Reder, 1988), 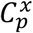 and 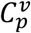 can be derived through the following expressions:

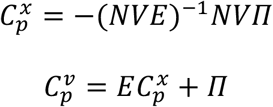

where *N* is the stoichiometric matrix, *V* is the diagonal matrix whose elements are the steadystate fluxes, *E* is the elasticity matrix with respect to metabolites and *Π* is the matrix of elasticities with respect to parameters.

Hence, flux and concentration control coefficients are computed for each reaction flux *i* and metabolite concentration *j* with respect to the system parameter *k* as:

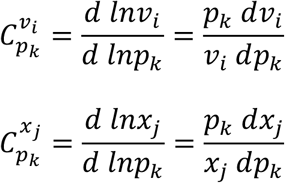

### Model description and calculation of control coefficients

The stoichiometric model that was used in this study (Hameri et al., 2019c) was systematically reduced from the *E. coli* iJO1366 genome-scale model (Orth et al., 2011) around the originally defined reaction subsystems of glycolysis, pentose phosphate pathway (PPP), tricarboxylic acid (TCA) cycle, glyoxylate cycle, pyruvate metabolism and the electron transport chain (ETC), and describes the aerobically grown physiology of *E. coli* (Supplementary Table S3). The reduction was performed through the redGEM and the lumpGEM algorithms (Ataman et al., 2017; Ataman and Hatzimanikatis, 2017), thus ensuring preservation of as much information as possible as well as that thermodynamic feasibility constraints are respected. This model constitutes of 337 metabolites participating in 647 reactions, which are in turn associated with 271 enzymes that serve as parameters in the NRA formulation. The model was curated with thermodynamic feasibility constraints using TFA (Henry et al., 2007; Salvy et al., 2019) and relevant fluxomics data (McCloskey et al., 2014). The representative steady state profiles of the metabolite concentrations and metabolic fluxes were chosen with Principal Component Analysis (PCA) as detailed in (Hameri et al., 2019b). Then, the populations of control coefficients were built using the ORACLE workflow (Andreozzi et al., 2016a; Miskovic et al., 2017; Miskovic and Hatzimanikatis, 2010; Tokic et al., 2020).

The CCs of the analyzed quantities (glycose uptake, pyruvate production, yield of pyruvate from glucose) with respect to the lumped reactions, exchange reactions, individual biomass building block contributions, and moieties were not considered in any study (Supplementary Table S3).

### Addressing variability in control coefficients

A common issue in MCA and in kinetic modeling is the uncertainty stemming from the scarcity of knowledge concerning the kinetic properties of enzymes (Miskovic and Hatzimanikatis, 2011; Miskovic et al., 2015; Miskovic et al., 2019b; Wang et al., 2004). The usual approach in addressing this issue involves the generation of a population of the CCs, and statistical analysis thereof. To form the NRA models, we need to select sets of CCs that will be representative of the generated population.

To select a representative set of CCs for our analysis, we took the population of 50’000 sets of FCCs and CCCs computed with ORACLE for the aerobically grown *E. coli* in (Hameri et al., 2019c). We first identified the vector of FCCs that was closest to the mean of the FCC distribution with respect to glucose uptake and selected it as the representative set. Four glucose uptake reactions in the model of *E. coli* exist with GLCptspp being responsible for 91.21% of the total flux through these reactions. We enforced this ratio in all performed NRA studies.

Since the model is constrained to grow on minimal media with glucose as its sole carbon source, the choice of the representative set will have a strong impact on the design criteria we wish to explore. To investigate the variability in results that this choice can induce, we additionally selected several “extreme” CC-sets through the use of PCA. We used nine principal components to describe the space of CCs with respect to glucose uptake, which lead to a coverage of 96.63% of the space variance. We selected the minimum and maximum corresponding CC-sets for each component (2 x 9), leading to a total of 19 sets. We then constructed 19 NRA models with these CC-sets and used them in the performed studies.

### Confidence Intervals and Bonferroni correction

For the computation of confidence intervals in Figure 2, we have used the Bonferroni correction in order to account for the multivariate nature of our study. In univariate studies, to account for the variability in samples, confidence intervals that contain the population mean with the probability 1 – α (typically, α = 5%) are added around each sample mean (Hameri et al., 2019a). However, univariate confidence intervals cannot be used when multivariate problems are studied, instead the Bonferroni’s correction of confidence intervals is frequently applied. In Bonferroni’s correction, for a problem with *p* variables, to ensure the level 1 – α for all variables simultaneously, we need to choose level 1 – α/*p* for each of individual variables. For instance, if we want to form confidence intervals for 10 variables with an overall 95% confidence level, then we need to use individual 99.5% confidence intervals.

### Thermodynamic constraints

To integrate thermodynamic constraints, we assumed that reactions operate in the directionality determined by the computed reference steady state. Thus, the concentrations of each metabolite in the respective cellular compartment need to be such as the 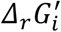 of each reaction remains negative. These constraints are written as a function of the standard Gibbs free energy change of the reaction 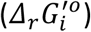 and the logarithmic concentrations of the participating metabolites, as introduced by (Henry et al., 2007). The 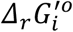 of each reaction is computed using the Group Contribution Method (Mavrovouniotis, 1990; Mavrovouniotis, 1991). These values are further adjusted to take into account the thermodynamic properties of the relevant cellular compartments; the pH gradient and electrochemical potential for transport reactions, and ionic strength of dissociated metabolites (Henry et al., 2006).

### Constraints on enzyme activities

Since the activity of an enzyme in the metabolic network could either be increased or decreased, but not both at the same time, we made use of integer variables in the formulation. Therefore, we split the catalytic activity deviations of our system, *E_k_*, into the continuous variables 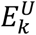 and 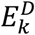, which denote the upregulation and downregulation of the gene encoding for enzyme *k*, respectively (Eqs. 11-14). As these should not have nonzero values simultaneously, we define the integer binary variables 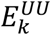 and 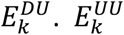 equals one if the gene catalyzing the enzyme *k* is upregulated and equals zero otherwise. In contrast, 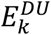 equals zero in the case of upregulation, and it is one for downregulation. As expressed in Eq. 12, only one of these variables can be active at a time, since deregulation cannot occur in both directions simultaneously, or they can both be inactive for the case of no change in the respective enzyme’s catalytic activity. To complete the formulation, these variables are further coupled to the above defined split enzymatic deviation variables through Eqs. 13 and 14. The integer binary variable *z_k_* is equal to zero if the activity of enzyme k is modified in the solution, and it equals to one otherwise (Eq. 11). ξ is a constant selected to be larger than the absolute value of the largest enzymatic deviation constraints, *lb_E_* and *ub_E_*, defined in Eq. 10.

### Software and optimization parameters

The computations were made on a Mac Pro workstation running Mac OS X version 10.11.6, equipped with a 2.7 GHz 12-Core Intel Xeon E5 processor and 32GB DDR3 memory, using MATLAB version R2016a and the IBM CPLEX solver version 12.5.1. Time limits for the solver were set as following: in Figure 2(a), for 2-fold (blue line) to 10 minutes, for 5-fold (orange line) to 30 minutes, and for 10-fold (yellow line) to 3 hours; in Figure 2(b), for all cases to 10 mins; in Figure 3, for all cases to 30 minutes; in Table 2, for all cases to 30 minutes; in the pyruvate case study (Figures 4–7), for all cases 3 hours.

## Conclusions

The NRA framework enables the consistent and sophisticated design of metabolic engineering strategies using MCA-based control coefficients. NRA is computationally faster and simpler than other approaches since the derivation of control coefficients does not require the numerical integration of non-linear kinetic models, and offers the implementation of a wide variety of metabolic engineering criteria. To our knowledge, this type of approach has never been applied to large or genome scale kinetic models of metabolism. Using a previously published large-scale kinetic model of *E. coli,* we demonstrated that the NRA formulation can be applied to large-scale metabolic networks. We used the PCA method to select a number of representative sets of kinetic parameters among their population, in order to effectively represent the uncertainty and flexibility of the kinetic model in respect to parametrization. One of the main advantages of NRA is that, being a constraint-based modeling method, it can accommodate the integration of biologically relevant bounds and constraints, which ensure that the proposed strategies are consistent with the entire system capabilities and limitations thereof. Since the NRA model predictions can be sensitive to the user-defined bounds on the allowable reaction flux, metabolite concentration and enzymatic expression deviations, the importance of including relevant physiological constraints, such as thermodynamic feasibility constraints, was discussed extensively. Focusing on the case of pyruvate production, a compound of great industrial interest, viable metabolic engineering strategies were shown to be readily derived using this formulation. Alternative solutions could also be generated and evaluated on their efficiency and potential implementation. We believe that this formulation will provide a refined alternative to computational genetic design, due to its simplicity and modularity, and that it will continue to be enhanced through the introduction of ever-growing omics data, and additional specialized constraints and objectives.

## Acknowledgements

S.T. and T.H. were supported by the Swiss National Science Foundation grant [315230_163423]. M.A. was supported by the RTD grant MicroscapesX within SystemsX.ch, the Swiss Initiative for Systems Biology evaluated by the Swiss National Science Foundation, and RobustYeast within ERA net project via SystemsX.ch. L.M. and V.H. were supported by the Ecole Polytechnique Fédérale de Lausanne (EPFL).

## Supplementary Material

Table S1: Metabolite concentration violation magnitudes for designs with two, four, and seven gene manipulations.

Table S2: List of the 51 generated alternative designs with the corresponding manipulations and magnitudes of manipulations, pyruvate productivity, and yield.

Table S3: List of aerobically grown *E.coli* model reactions, metabolites, and parameters considered in the study.

Figure S1: Absolute differences of fluxes in the network for the Alternative 1 design (Group 1). Blue/pink arrows and numbers denote an up-/down-regulation of the genes encoding for the respective enzyme and the corresponding fold-change value.

Figure S2: Absolute differences of fluxes in the network for the Alternative 25 design (Group 2). Blue/pink arrows and numbers denote an up-/down-regulation of the genes encoding for the respective enzyme and the corresponding fold-change value.

Figure S3: Absolute differences of fluxes in the network for the Alternative 51 design (Group 3). Blue/pink arrows and numbers denote an up-/down-regulation of the genes encoding for the respective enzyme and the corresponding fold-change value.

Figure S4: Absolute differences of fluxes in the network for the Alternative 40 design (Group 4). Blue/pink arrows and numbers denote an up-/down-regulation of the genes encoding for the respective enzyme and the corresponding fold-change value.

Figure S5: Absolute differences of fluxes in the network for the Alternative 45 design (Group 5). Blue/pink arrows and numbers denote an up-/down-regulation of the genes encoding for the respective enzyme and the corresponding fold-change value.

