## Supplementary figures and images for "Constraint-based metabolic control analysis for rational strain engineering"

### Supplementary figure S1

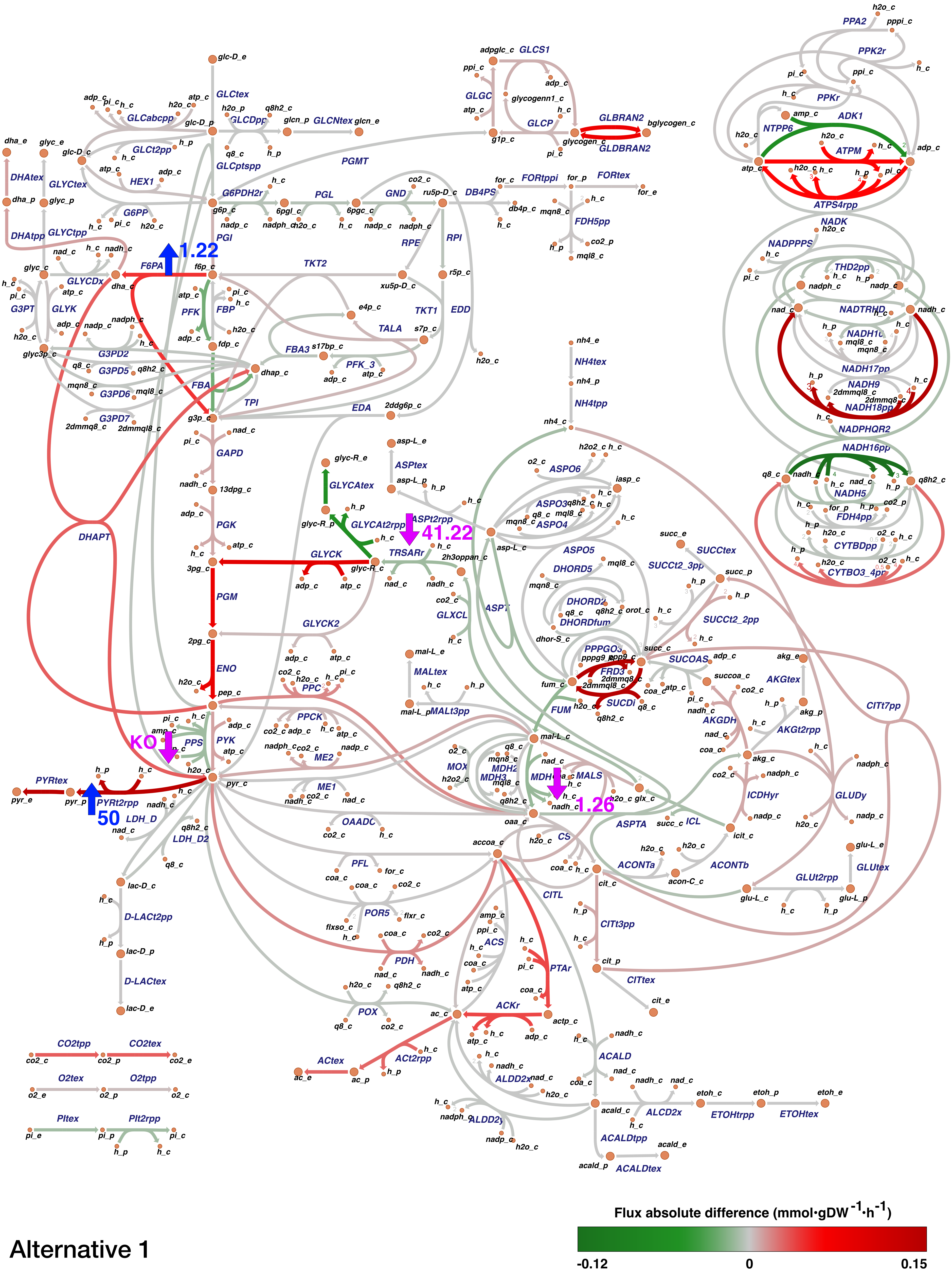


Alternative 1

### Supplementary figure S2

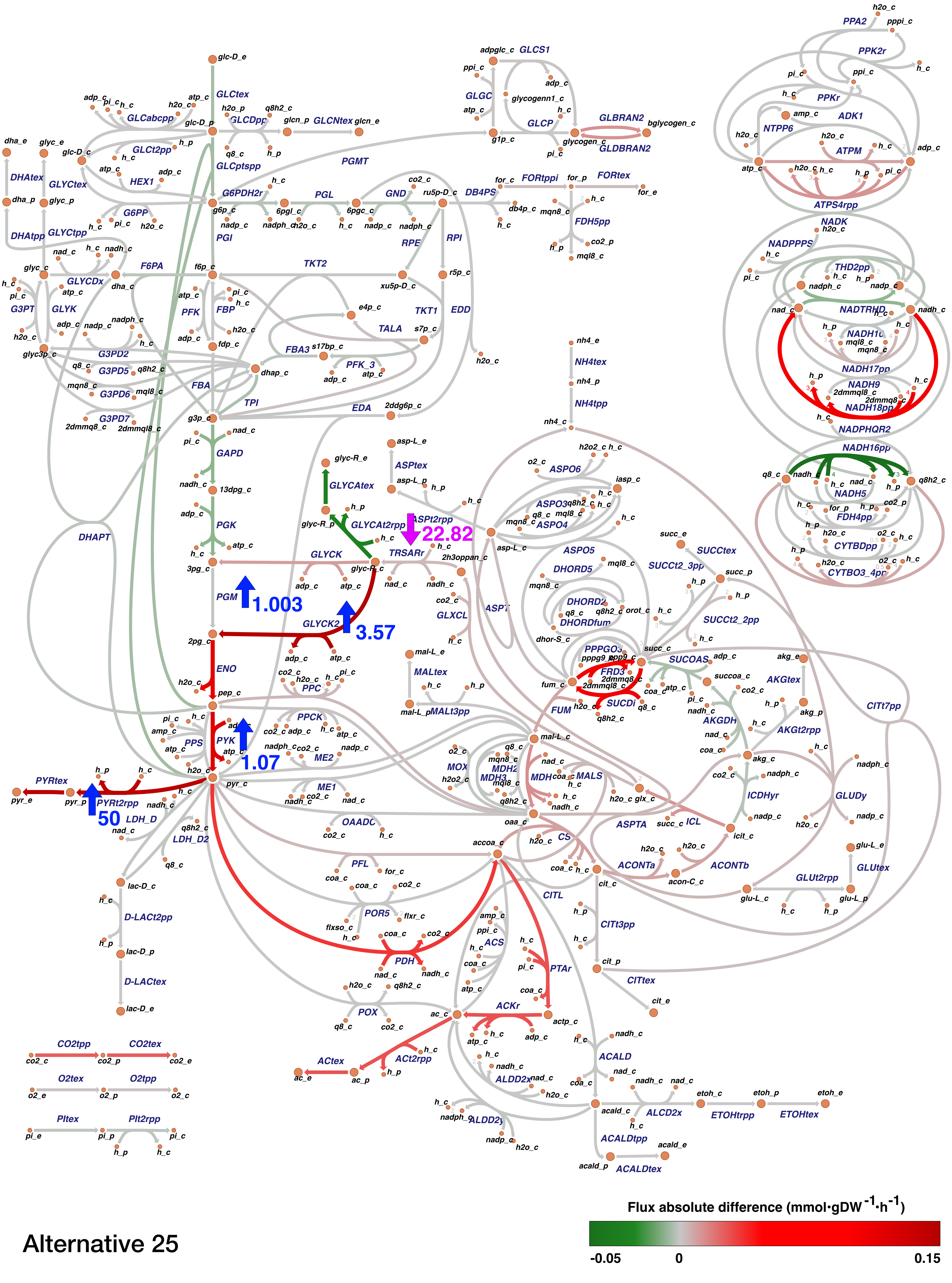


Alternative 25

### Supplementary figure S3

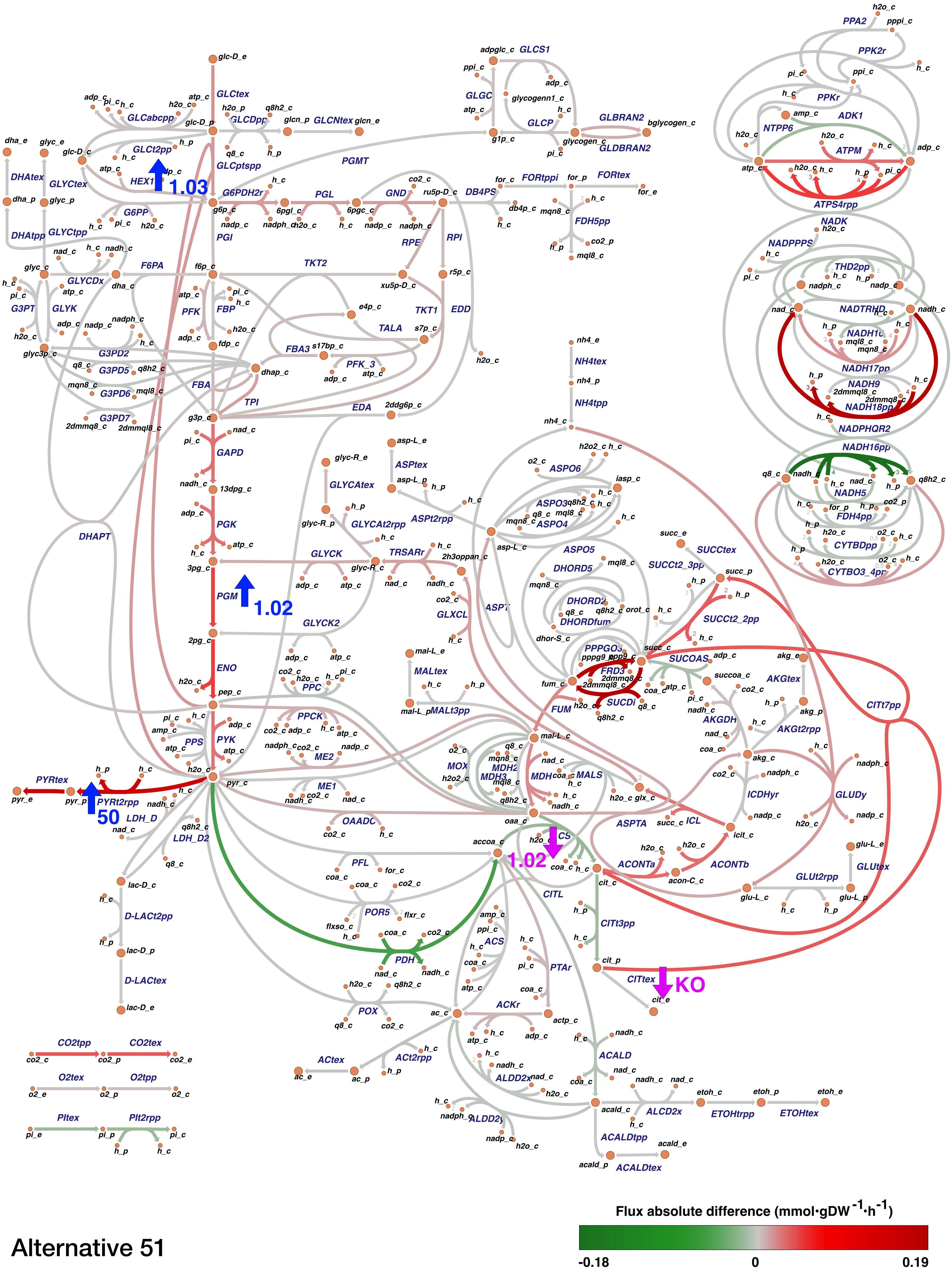


Alternative 51

### Supplementary figure S4

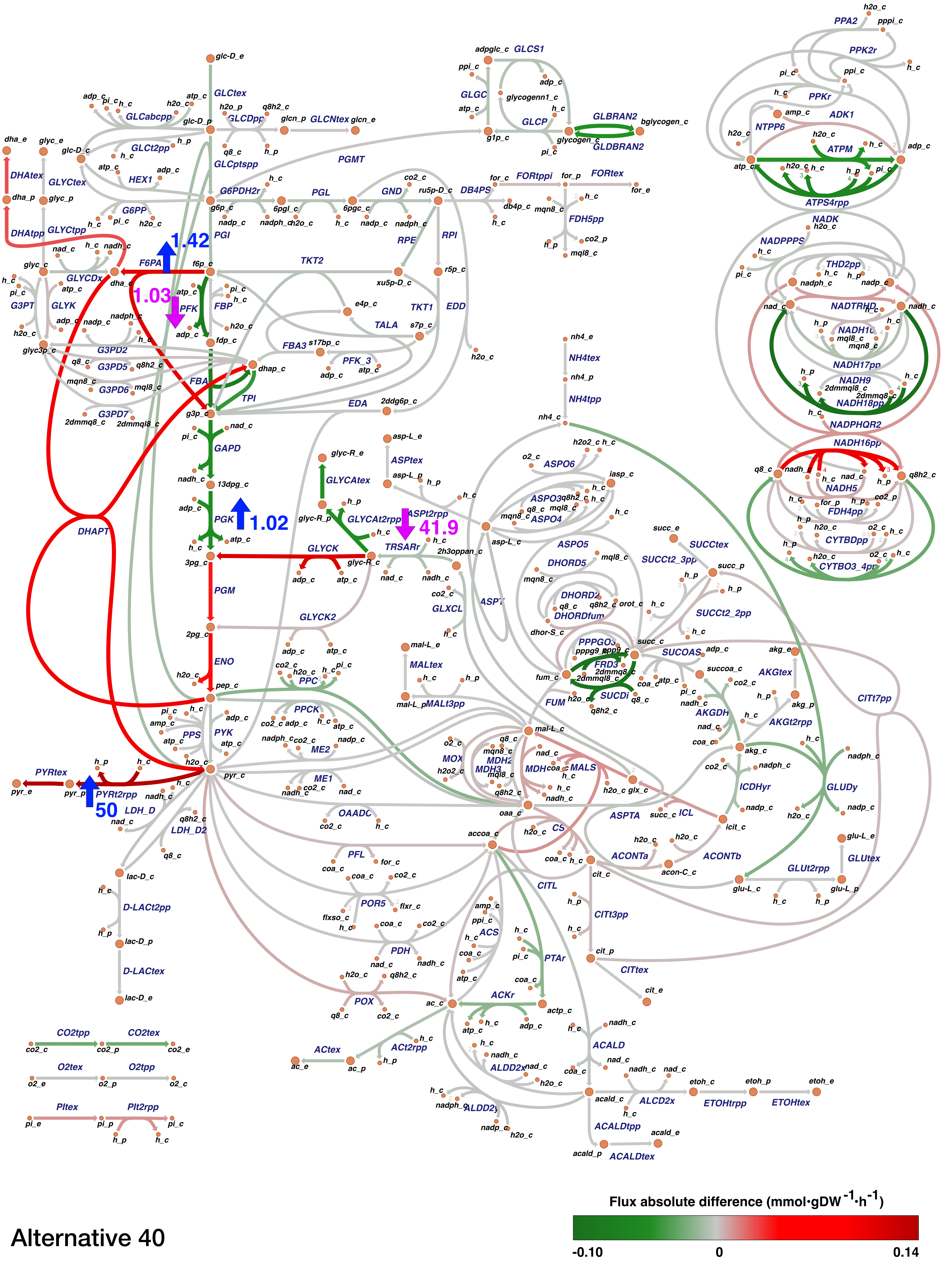


Alternative 40

### Supplementary figure S5

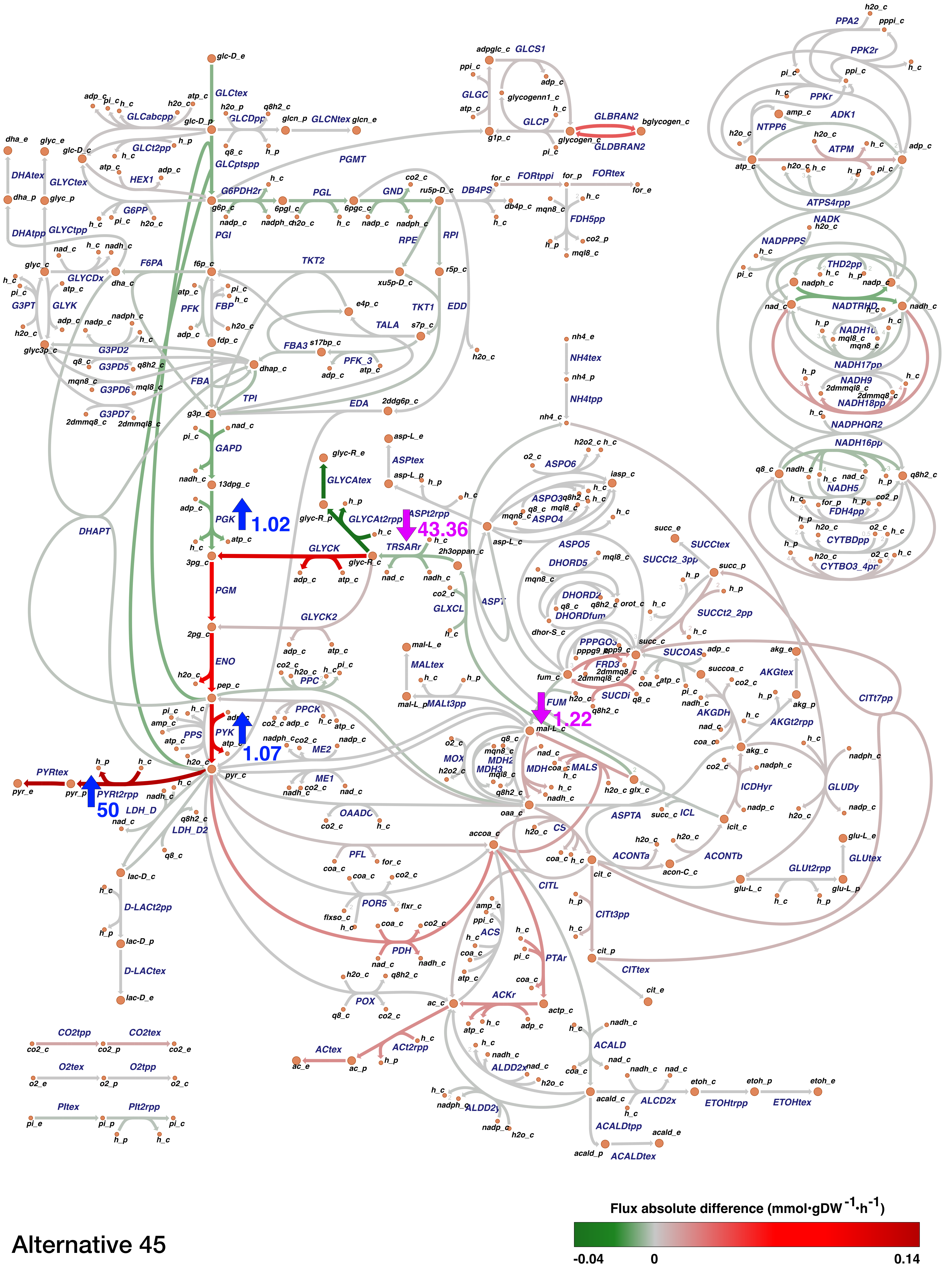


Alternative 45
